# Extracellular Spike Waveform and Receptive Field Classes Capture Partially Independent Structure in Awake Macaque LGN

**DOI:** 10.1101/2023.11.22.568065

**Authors:** Shi Hai Sun, Nathaniel J. Killian, Jae-Ik Lee, John S. Pezaris

## Abstract

Extracellular spike waveforms vary substantially across neurons, but whether waveform-defined classes in the macaque LGN correspond to differences in receptive field organization and functional feature space remains unclear. Here, we characterized extracellular spike waveforms and receptive field properties in the LGN of awake monkeys using multi-electrode recordings during the presentation of colored noise visual stimuli. Extracellular spike waveforms were classified using a PCA-based graph clustering workflow, which identified distinct waveform classes that differed in spike shape statistics. Receptive fields (RFs) were classified by cone-response properties, and we found that waveform-defined classes were associated with structured differences in LGN feature space. Waveform classes differed in their proportions of RF classes and showed differences across RF properties and spiking characteristics. Multivariate analyses showed that both waveform and RF class were associated with position in feature space, and that waveform class explained variance beyond that accounted for by RF label, indicating that the two schemes captured partially independent structure. To test whether this structure generalized to unseen units, we trained cross-validated classifiers to predict waveform and RF class from multivariate physiological measurements. Both labels were recovered above permutation-based chance levels, although confusion between classes indicated substantial overlap rather than complete separation. These findings present a survey of extracellular spike waveform classes in the awake macaque LGN and suggest that waveform labels capture meaningful structure related to RF organization and visual response properties.

**Key Points:**

- Electrodes placed near brain cells record electrical signals that are usually assumed to come from the main body of a single nerve cell.
- In the lateral geniculate nucleus, the relay between the eye and the visual cortex, these signals vary in shape, but whether shape relates to visual responses is unknown.
- In awake monkeys, we found four signal shapes, including some that may come from nerve fibers rather than cell bodies.
- Signal shape and the cells’ color responses were related, but each carried information the other did not.
- Signal type should therefore be considered when interpreting electrode recordings of the visual system in health and disease.

## Introduction

The lateral geniculate nucleus (LGN) of the thalamus serves as an important part of the early stages of visual information processing in mammals, acting as the primary relay station between the retina and the visual cortex. In primates, LGN neurons are commonly described by distinct visual response profiles, including luminance-sensitive, color-opponent, and short-wavelength-cone-sensitive responses (Sherman & Guillery, 2006; Solomon & Lennie, 2007). These response profiles are related to the classically described magnocellular, parvocellular, and koniocellular pathways, which typically differ in temporal dynamics, chromatic sensitivity, spatial resolution, and cone inputs (Wiesel & Hubel, 1966; Schiller & Malpeli, 1978; Kaplan & Shapley, 1982; Derrington & Lennie, 1984; Hubel & Livingstone, 1990; Maunsell *et al*., 1999; Reid & Shapley, 2002; Tailby *et al*., 2008).

Much of our understanding of LGN function stems from electrophysiological recordings of extracellular spikes. Typically, these signals are biphasic with a dominantly negative voltage excursion, allowing for tentative classification of neuronal cell types: broader waveforms are often associated with excitatory neurons, while narrower waveforms are often associated with inhibitory neurons (Henze *et al*., 2000; Barthó *et al*., 2004). However, recent advances in dense multi-electrode arrays and spike-sorting algorithms have revealed a wider range of waveform shapes across species and brain regions. These waveform variations include triphasic waveforms in rat hippocampus (Barry, 2015) and superior colliculus (Sibille *et al*., 2022), and positive-spiking and long-broad negative waveforms in cat visual cortex (Gold *et al*., 2009; Sun *et al*., 2021) and human cortex (Paulk *et al*., 2022). Many of these nontraditional waveforms are thought to reflect non-somatic activity, axonal signals, dendritic return currents, or intermediate waveform shapes; revealing a broader diversity of extracellular waveform types than previously appreciated (reviewed in <u>Sibille *et al*., 2026; Sun & Ibbotson, 2026; Siegle & Steinmetz, 2026</u>). Such findings challenge the conventional view of extracellular spikes as simply somatic signals and underscore the limitations of electrode sampling biases (Olshausen & Field, 2005; Talebi & Baker, 2016).

Although extracellular waveform shape contains information related to neuronal identity or functional properties in many brain regions (Henze *et al*., 2000; Barthó *et al*., 2004; Sun *et al*., 2021; Lee *et al*., 2021, 2026; Carr *et al*., 2026), the functional relevance of waveform diversity in the macaque LGN remains unclear. Specifically, it is not known whether the cellular sources in LGN that give rise to different waveforms, such as distinct cell types or non-somatic compartments, also differ in receptive field structure, cone-response properties, or spike-train statistics. This question is well suited to graph-based approaches that organize extracellular waveform features into interpretable low-dimensional spaces (Lee *et al*., 2021, 2026), which were built on methods developed for single-cell analysis, including Seurat-based graph construction and clustering (Hao *et al*., 2024), and provide a way to test whether waveform-defined classes occupy distinct regions of functional feature space (i.e., the combined set of spike waveform, receptive field, cone-response, and spike-train measurements used to describe each unit), and whether they do so independently of RF identity.

Here, we characterize extracellular spike waveform diversity in the LGN of awake rhesus macaques and ask whether waveform-defined classes are associated with distinct regions of functional feature space. We focus on receptive-field-mapped units with reliable spike waveforms and describe receptive fields by their measured cone-response properties as LM, LM-opposing, or S, which reflect functional response profiles related to, but not equivalent to, magnocellular, parvocellular, and koniocellular LGN pathways, respectively. We then use graph-based clustering of waveform principal components to define extracellular waveform classes and examine how these classes relate to spike shape statistics, receptive field properties, cone-response labels, and spike-train characteristics. Since waveform class and cone-response label are themselves associated, we further ask whether waveform class accounts for structure in feature space that is not already explained by RF label. This approach allows us to test whether extracellular waveform labels capture meaningful structure in the functional organization of the awake primate LGN.

## Methods

### Ethics

Recordings were made from three awake adult rhesus monkeys (*Macaca mulatta*, 3 males, 19–20 kg), sourced from Alpha Genesis Primate Research Center. The animals were maintained in the AAALAC (Association for Assessment and Accreditation of Laboratory Animal Care) accredited animal facility at the Massachusetts General Hospital. All research procedures were approved by the Massachusetts General Hospital Institutional Animal Care and Use Committee (IACUC) and were carried out according to the NIH Guide for the Care and Use of Laboratory Animals and the ARRIVE guidelines.

### Animal preparation

The animals were surgically implanted with custom titanium head-holding posts and recording cylinders that allowed bilateral access to the LGN. Animals were trained to sit in a primate chair (B & M Plastics, Inc.), and were placed in a shielded recording chamber (Crist Instrument Co.) during recordings. The animals were positioned so that their eyes were 43 cm from a 22-inch CRT monitor (Viewsonic P220f) with neutral gaze position near the center of the monitor.

Gaze location was monitored by an infrared video camera at a 500 Hz sampling rate (High-Speed Primate, SensoMotoric Instruments GmbH), with infrared illuminators stacked to yield a single corneal reflection. Software (iView X, SensoMotoric Instruments) was used to adjust the gains and offsets of the pupil-corneal reflection distance. Gaze position signals were then sent as analog signals to a behavioral control computer, where additional calibration was performed using a two-dimensional quadratic fit to nine calibration points.

### Behavioral task

A trial-based task was used to map the receptive fields of neurons (Figure 1). To begin each trial, the animal was required to fixate on a centrally presented small, circular target (0.2° diameter white spot) for 400 ms. The visual mapping stimulus then began with the fixation point overlaid on the animated noise so that it continued to be visible, and the animal was required to continue maintaining fixation within a circular window (2° diameter; typical fixation was substantially tighter than this window; see Figure S4) for the duration of the stimulus (2.5–5.0 seconds, mode of 2.5 seconds; typical fixation performance was substantially tighter than this window). Successful fixations through the entire stimulus were rewarded with drops of sweet liquid and pleasant audio feedback. Data from successful trials were used for mapping analysis (4–215 trials per experiment, median of 31 trials), whereas data from all trials were used for spike waveform analysis.

**Figure 1.**
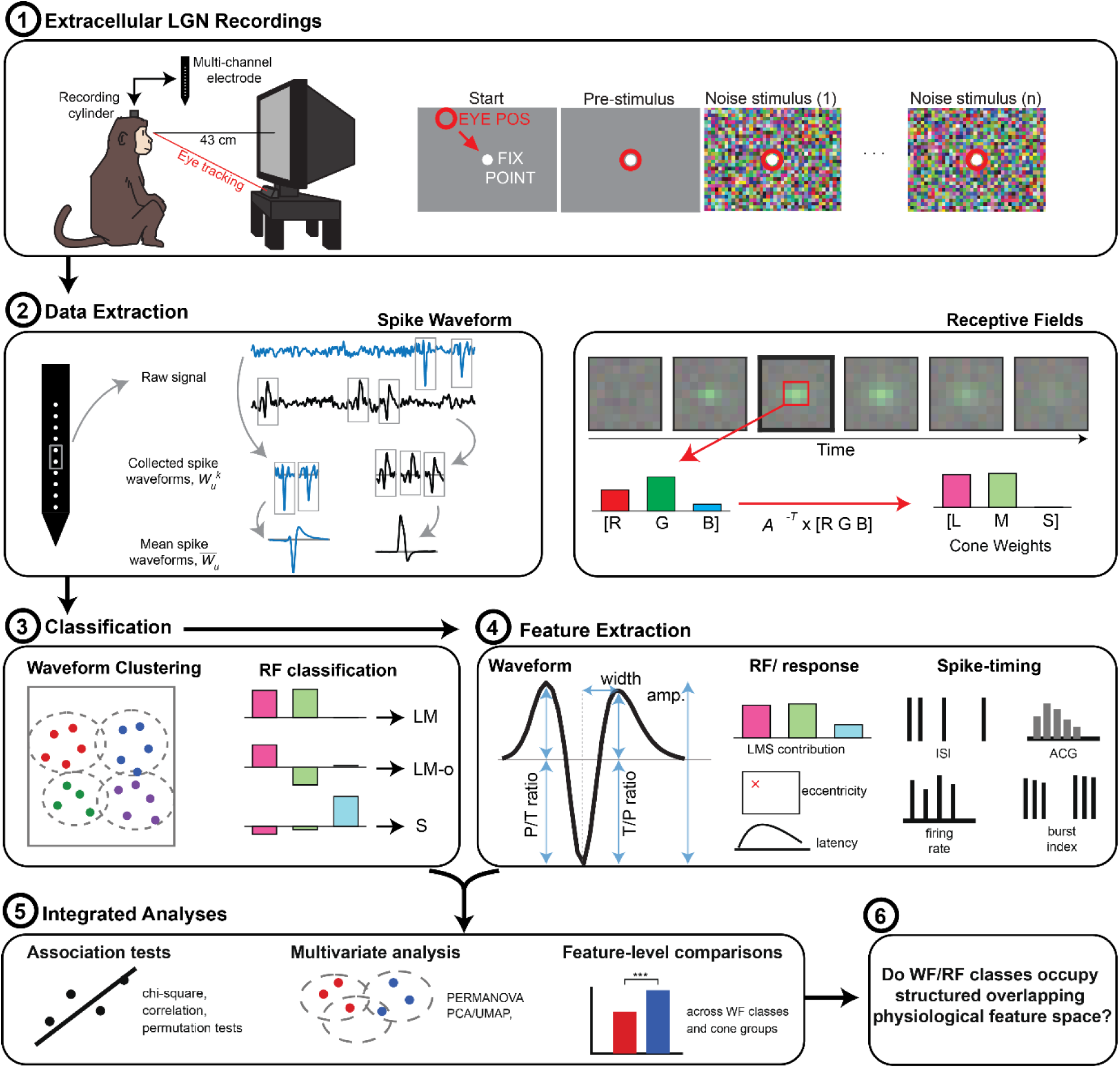
Experimental design and analysis workflow. **(1)** Awake rhesus macaques maintained fixation on a central target while multichannel electrodes recorded extracellular activity from the lateral geniculate nucleus (LGN). Eye position was monitored continuously during presentation of colored noise stimuli. **(2)** Individual spike waveforms were extracted from the raw voltage signal and averaged to obtain a mean extracellular waveform for each unit. Chromospatiotemporal receptive fields (RFs) were estimated by spike-triggered averaging of the colored noise stimulus. RGB responses were transformed into long-, middle-, and short-wavelength cone weights. **(3)** Mean waveforms were classified using a principal-component- and graph-based clustering workflow. RFs were classified as LM, LM-opposing (LM-o), or S according to their measured cone-response structure. **(4)** Waveform-shape, RF-response, and spike-timing features were extracted for each unit. **(5)** Relationships between waveform and RF classes were evaluated using categorical association tests, multivariate feature-space analyses, and feature-level comparisons. **(6)** These analyses tested whether waveform and RF classes occupied structured and overlapping regions of physiological feature space. WF, waveform; RF, receptive field; ISI, interspike interval; ACG, autocorrelogram.

### Visual Stimuli

Custom software (PLECS, Pezaris Lab Experiment Control System) was used to display noise stimuli that spanned both luminance and chrominance spaces, and to control behavioral state and data collection (Killian *et al*., 2016). Visual stimuli were presented on a CRT display (Viewsonic P220f) at a 160 Hz refresh rate with 8 bits per color channel. The display area measured 40 by 30 cm (400 by 300 pixels), subtending 53° by 40° of visual angle. The gamma look-up table of the display was linearized such that incremental changes in the driven level of the red, green, or blue guns produced approximately equal changes in spectral radiance. Spectral radiance values were measured using a PR650 spectrophotometer (Photo Research, Inc.).

Two classes of noise stimuli were presented. High-resolution stimuli were 400 by 300 pixels with power spectra proportional to 1/*f^β^*, generated in the frequency domain by imposing the desired power spectrum on random phase and inverting the Fourier transform (<u>Simoncelli & Olshausen, 2001</u>). Low-resolution stimuli were 80 by 60 pixels with a white power spectrum (*β* = 0). The high-resolution stimuli used *β* values of 1.5, 2.5, 3.14, 3.5, or 4.0, where higher values yield more naturalistic spectra with greater spatial and temporal correlations. Each recording used a single mapping stimulus. Example stimuli are shown in Figure S1.

### Electrode implantation

LGN recordings were made with acutely inserted electrodes (*n* = 98 recordings; 77 multichannel recordings, 21 single-channel recordings; three animals) or chronically implanted electrodes (*n* = 10 recordings; one animal). Acute electrodes were either a linear 16-electrode array with 200 µm channel spacing (Plexon V-Probe), a linear 16-electrode tetrode configuration array (four groups of four contacts) with 50 µm intra-group spacing and 450 µm inter-group spacing (Plexon U-Probe), custom single linear 16-electrode array with varying channel spacing (50–300 µm), or a single traditional tungsten electrode (FHC, Inc.). Chronic electrodes were custom 64-channel microwire bundles designed to splay at depth.

Neural responses were recorded by a data acquisition system (Power 1401, Cambridge Electronic Design) and software (Spike 2, Cambridge Electronic Design) that sampled at 40 kHz for all electrodes and channels, to 16-bit resolution. Single-channel units recorded with traditional tungsten electrodes were included in waveform clustering (see Waveform clustering) and in the class-composition analyses of Figure 3, but were excluded from all receptive-field feature analyses, which used multichannel units only.

### Spike sorting

Extracellular signals were automatically sorted using KiloSort 2.5 (Pachitariu *et al*., 2016), and the output was manually curated using phy (Rossant *et al*., 2016). During manual curation, clusters were cleaned by drawing boundaries in principal component space to remove abnormal waveforms and were then assigned as either *good* units or *noise*. *Good* units showed clear isolation from other clusters in principal component space and evidence of a refractory period in the autocorrelogram, defined by a central dip to approximately zero for at least 1 ms. Clusters were merged when they had similar waveforms, overlapping principal component distributions, shared refractory structure in the cross-correlogram, and/or evidence of drift across adjacent channels. *Noise* clusters were excluded from subsequent analyses. Spike sorting examples are shown in Figure S2.

### Extracellular spike waveform analysis

Each single unit was represented by its mean extracellular spike waveform, *W̅_u_*. For each unit, up to 2,000 spike waveforms were randomly selected from the identified spike times. If fewer than 2,000 spikes were available, all spikes were used. Waveforms were extracted in 4.5 ms windows spanning 60 samples before and 120 samples after the spike time at a 40 kHz sampling rate, and were averaged to obtain *W̅_u_*.

For multi-channel recordings, the channel with the largest peak-to-peak mean waveform amplitude was selected for waveform analysis. Waveform quality was quantified using a signal-to-noise ratio (SNR), defined as the peak-to-peak amplitude of the mean waveform divided by the standard deviation of residual waveform noise after subtracting the mean waveform from each individual waveform. Only RF-mapped units recorded on multi-channel electrodes with waveform SNR greater than 3 were included in the waveform clustering and feature-space analyses, yielding 226 units. The median waveform SNR of the included population was 5.14.

#### Spike waveform classification

Mean spike waveforms, *W̅_u_*, were classified using a graph-based clustering workflow derived from PhysMAP and implemented with Seurat (Stuart *et al*., 2019; Hao *et al*., 2024; Lee *et al*., 2026). Before clustering, waveforms were smoothed with a three-sample moving average, baseline-subtracted using the first and last 10 samples, and normalized by their maximum absolute amplitude. Processed waveforms were imported into Seurat as a waveform assay, centered, scaled, and reduced by principal component analysis. Based on inspection of the variance explained plot, 20 waveform PCs were used for graph construction and clustering.

A shared nearest-neighbor graph was constructed from the selected PCs, and waveform classes were identified using Louvain community detection (Blondel *et al*., 2008). Clustering resolution was selected by testing resolutions from 0.1 to 2.0 across 100 bootstrap iterations, each using 80% of units sampled without replacement. The final resolution was the last value for which the increase in graph modularity remained significant and exceeded 0.01; testing stopped at the next resolution when either criterion was no longer met. The variance explained and resolution optimization plots are shown in Figure S3. UMAP was calculated from the selected waveform PCs using cosine distance for visualization (McInnes *et al*., 2020). The resulting groups are referred to as waveform classes rather than cell types.

#### Spike waveform metrics

Quantitative metrics were extracted from each unit’s mean spike waveform, *W̅_u_*, after baseline subtraction.

Waveform amplitude was defined as the peak-to-peak range of the baseline-subtracted waveform (maximum minus minimum).

The polarity index was defined as:

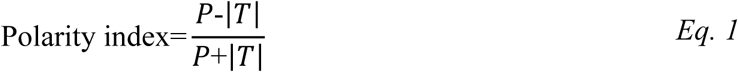

where *P* is the maximum positive deflection and |*T*| is the absolute amplitude of the negative trough, yielding values from −1 (purely trough-dominant) to +1 (purely peak-dominant).

Waveform duration was defined as the time from the trough to the following positive peak, where the following peak was identified as the largest positive local maximum occurring within 1 ms after the trough. When no positive local peak was detected within this window, duration was instead measured to the onset of the first sustained flat interval: a 0.2 ms window in which the amplitude range was less than 5% of |*T*| and all sample-to-sample changes were less than 5% of the maximum post-trough step. When neither condition was met, duration was measured to the end of the recording window.

The pre-peak to trough ratio (PT-ratio) was defined as the amplitude of the largest positive value within 1 ms preceding the trough, divided by |*T*|. The trough to post-peak ratio (TP-ratio) was defined as the absolute amplitude of the following positive peak divided by |*T*|. When duration was determined by flattening rather than a detected peak, TP-ratio was set to zero.

Because duration, PT-ratio, and TP-ratio are referenced to the trough, these metrics were not calculated for positive-dominant waveforms and were excluded from the corresponding analyses. Waveform amplitude and polarity index were retained for all units.

### Receptive field analysis

#### Spike-triggered averaging

Receptive fields were estimated from successful fixation trials using spike-triggered averaging of the colored noise stimulus. For each single unit, the spike-triggered average (STA; Schwartz *et al*., 2006) was computed by averaging the stimulus frames preceding and following each spike:

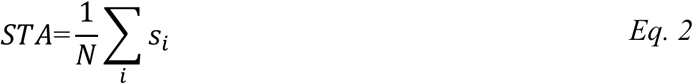

where *s_i_* is the stimulus segment surrounding the *i*^th^ spike and *N* is the total number of spikes used for the receptive field estimate. The STA was computed over a temporal window spanning 25 stimulus frames before to 10 frames after each spike, corresponding to time lags of −156.25 ms to 62.5 ms in steps of 6.25 ms. To correct for correlations in the colored noise stimulus, the STA was whitened by multiplying it by the inverse of the stimulus covariance matrix (Sharpee, 2013). The resulting chromospatiotemporal receptive field contained time, horizontal position, vertical position, and color dimensions, with color represented by red, green, and blue monitor phosphor values.

#### RF position and latency

At each spatial and temporal location, the magnitude of the RGB receptive field vector was calculated as:

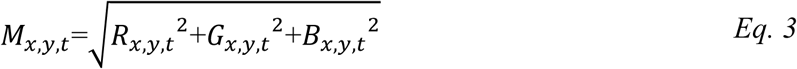

where (R), (G), and (B) are the receptive field values for the red, green, and blue phosphor channels. The spatiotemporal RF peak was defined as:

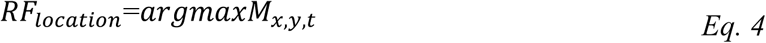

RF position was defined by the horizontal and vertical coordinates of the spatiotemporal peak, and RF latency by the corresponding temporal lag.

#### RF spatial window

For RFs mapped with high-resolution stimuli, the RGB vector magnitude at the peak temporal frame was fitted with an oriented two-dimensional Gaussian within a 3° region centered on the RF. Horizontal and vertical extents were taken as the full widths at 60% of maximum, converted to degrees of visual angle, and corrected for eccentricity; their mean defined a square spatial window centered on the RF peak. For RFs mapped with low-resolution stimuli, the peak pixel was used instead.

#### RF temporal window

The temporal RF response was obtained by averaging the receptive field within the spatial window at each time lag, and a two-component Gaussian was fitted around the peak frame (polarity inverted for OFF-dominant RFs). Frames were included when the response matched the RF peak polarity and exceeded the noise level by at least one standard deviation.

#### RF strength

RF strength was quantified from the RGB magnitude within the selected spatial and temporal windows. For each included pixel, the magnitude of the RGB vector magnitude was calculated using Equation 3, and RF amplitude was defined as the mean magnitude across all selected pixels:

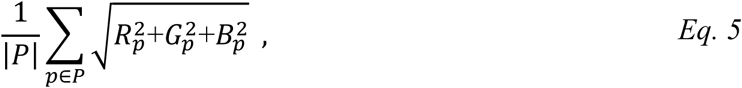

where P is the set of pixels within the RF spatial and temporal windows, and |P| is its size.

A noise distribution was constructed from the 10 acausal RF frames at positive post-spike lags by repeatedly sampling pixel groups matched in size to the RF window (see Supplementary Methods). RF strength was expressed as a *z*-score:

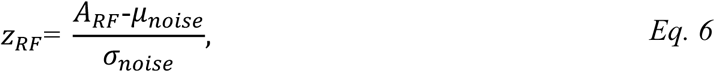

where *μ_noise_* and *σ_noise_* are the mean and standard deviation of the resulting noise distribution. The RF *z*-score was used to quantify receptive field response strength and to identify units with measurable receptive fields.

#### Cone weights calculation

RF color responses were converted from monitor-phosphor coordinates to cone coordinates using the method of <u>Horwitz & Albright, (2005)</u>. Before cone conversion, each RGB channel of the STA was baseline-corrected by subtracting its mean over the acausal frames (lags ≥ 0 ms). The RGB temporal response was then averaged within the RF spatial window and transformed into long-, middle-, and short-wavelength cone coordinates:

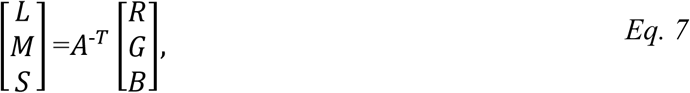

where *A*^-^*^T^* is the inverse transpose of the 3 × 3 RGB-to-LMS transformation matrix. A session-appropriate transformation matrix was selected according to the monitor calibration and shielding configuration used during each recording. These matrices were derived from pairwise inner products between the measured spectral radiance of each CRT phosphor and the Stockman–Sharpe 10° cone fundamentals (Stockman & Sharpe, 2000). Spectral radiance was measured with a PR650 SpectraScan colorimeter (Photo Research, Inc.).

Cone weights were calculated over the selected temporal RF frames and normalized by the sum of their absolute values:

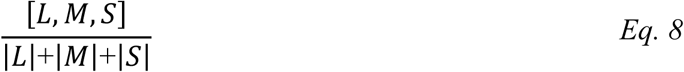

##### Cone weight noise

Noise distributions for the L-, M-, and S-cone weights were estimated from the same acausal RF frames used to calculate the RF *z*-score. Acausal RGB values were transformed into cone coordinates using Equation 7, and the standard deviation of each resulting cone-weight distribution was used as the corresponding cone-specific noise level.

##### RF classification

Mapped RFs were classified according to their normalized L-, M-, and S-cone weights. Classification was performed sequentially. RFs were first labeled LM-opposing when the L-and M-cone weights had opposite signs and the absolute magnitudes of both weights exceeded two standard deviations of their corresponding cone-noise distributions. Among the remaining RFs, RFs were labeled S when the S-cone weight either had the largest absolute magnitude of the three cone weights or exceeded three standard deviations of the S-cone noise distribution. All remaining RFs were labeled LM and were characterized primarily by same-sign L- and M-cone contributions.

These labels describe the measured cone-response structure of each RF and do not assign anatomical magnocellular, parvocellular, or koniocellular identity. LM, LM-opposing, and S response profiles are functionally related to classical magnocellular, parvocellular, and koniocellular response patterns, respectively, but are not treated as equivalent to those anatomical pathways here.

To summarize cone-response structure as continuous scalar features, two indices were derived from the normalized cone weights. The LM index quantified the balance and sign relationship between the L- and M-cone weights and was computed as

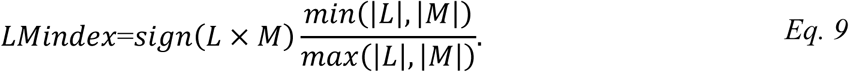

Values near +1 indicate balanced same-sign L- and M-cone contributions, values near –1 indicate balanced opponent L- and M-cone contributions, and values near 0 indicate dominance by one cone or weak contribution from one cone. The S index was defined as the absolute normalized S-cone weight,

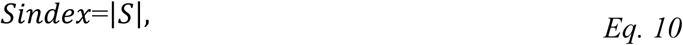

with higher values indicating greater S-cone contribution.

### Spike-train metrics

#### Spike rate

Mean spike rate was calculated as the total number of spikes divided by the duration of the analyzed recording period.

#### Burst index

Bursting was quantified using the classical LGN burst definition. A burst began when an interspike interval greater than 100 ms was followed by two or more spikes separated by interspike intervals shorter than 4 ms (Sherman, 2001; Wang *et al*., 2006). The burst index was calculated as the proportion of all spikes that occurred within identified bursts.

#### Autocorrelogram metrics

A high-resolution autocorrelogram was calculated from all nonzero pairwise spike-time lags within ±20 ms using 0.1 ms bins. The zero-lag bin was removed, and the ACG was normalized by spike count and bin width. Before feature extraction, the ACG was smoothed using a 0.5 ms moving average.

ACG peak latency was defined as the positive lag of the maximum smoothed ACG value between 2 and 20 ms.

ACG refractory recovery was defined as the first positive lag between 0 and 10 ms at which the smoothed ACG reached 50% of its post-refractory baseline (the mean rate between 10 and 20 ms) and remained above it for at least 0.5 ms.

#### Local interspike-interval variability

Local spike-timing variability was quantified using the local coefficient of variation (CV_2,*i*_; <u>Holt *et al*., 1996</u>). For each pair of consecutive interspike intervals,

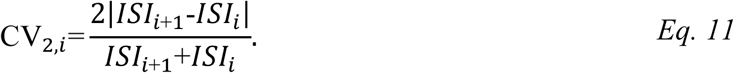

The mean CV_2,*i*_ across all valid adjacent interspike-interval pairs was used as the unit-level CV_2_ value. Larger values indicate greater local variability in spike timing.

### Statistical Analysis

Statistical analyses were performed in MATLAB, with significance defined as *p* < 0.05. All complete finite observations meeting the predefined inclusion and quality-control criteria were retained; observations were excluded only when a metric could not be calculated reliably or the recording failed quality control. No statistical outlier removal was applied.

Because stimulus statistics influence spike-train measurements (Almasi *et al*., 2022; Sanchez *et al*., 2023), all spike-train and feature-space analyses used only recordings obtained with the high-resolution 400 × 300 pixel stimulus.

Absolute anatomical depth and LGN layer boundaries could not be reconstructed reliably across recording sessions. Depth was therefore not included as a physiological feature in the primary analyses. An exploratory comparison was restricted to nine approximately linear penetrations for which relative within-penetration recording depth could be compared across RF classes. These relative measurements were used descriptively and were not treated as estimates of anatomical layer identity.

### Univariate comparisons

Differences among waveform or RF classes were assessed using one-way ANOVA for approximately normally distributed variables with homogeneous variances, Welch ANOVA for approximately normally distributed variables with unequal variances, and the Kruskal-Wallis test otherwise. Positive-dominant waveforms were excluded from analyses of waveform duration, PT-ratio, and TP-ratio, as described above. LM index and S index were not tested across RF classes, because those classes are defined by the cone weights from which both indices are derived.

Omnibus p-values were corrected across each family of metrics using the Benjamini–Hochberg false-discovery-rate procedure, and corrected values are reported throughout. Post hoc pairwise comparisons were performed following significant omnibus tests, using Tukey’s test after ANOVA and Dunn’s test after Kruskal-Wallis, with false-discovery-rate correction applied within each metric. Effect sizes were reported as η² for ANOVA and ε² for Kruskal–Wallis.

To assess the contribution of each metric to class separation while accounting for the other metrics, we additionally computed permutation feature importance from the cross-validated classifier described below: each metric was shuffled within the held-out fold and importance was defined as the resulting reduction in balanced accuracy, averaged over 20 cross-validation repeats.

Descriptive values in the Results are reported as medians unless otherwise noted; boxplots show medians and interquartile ranges.

### Multivariate feature-space analysis

Waveform and RF classifications were analyzed in a common feature space comprising RF eccentricity, RF latency, spike rate, burst index, CV_2_, ACG peak latency, ACG refractory recovery, and RF z-score. LM and S indices were excluded because the RF labels are defined by the cone weights from which they derive; retaining a shared feature set also allows the two classifications to be compared on identical terms.

Rows containing missing or nonfinite values were removed separately for each feature matrix, and each retained variable was standardized to zero mean and unit variance. No multivariate outlier removal was applied.

Principal component analysis was performed on each standardized feature matrix, and the first two principal components were used for visualization. Two-dimensional UMAP embeddings were generated using Euclidean distance, 15 nearest neighbors, a minimum distance of 0.1, and a fixed random initialization.

Overall differences among waveform or RF classes were evaluated using one-way permutational multivariate analysis of variance (PERMANOVA) based on Euclidean distances. Statistical significance was estimated using 10,000 permutations of the class labels. PERMANOVA effect size was reported as

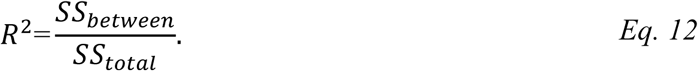

Pairwise PERMANOVA tests were performed between all class pairs with 10,000 permutations each and Benjamini–Hochberg correction within each classification. Since PERMANOVA is sensitive to differences in multivariate dispersion as well as location, homogeneity of dispersions was tested by permuting the distances of each unit to its group centroid (PERMDISP; Anderson, 2006), using 10,000 permutations.

Because waveform class and RF label are themselves associated, we fitted a two-way PERMANOVA to test whether each factor explained variance beyond the other. Each term was tested marginally, with the null generated by restricted permutation of that factor’s labels within levels of the other factor (Anderson & Braak, 2003), and effect size reported as partial R².

Class separability was additionally quantified by cross-validated classification. A regularized linear discriminant classifier was trained on the standardized features using stratified five-fold cross-validation, repeated 20 times. Performance was expressed as balanced accuracy (the mean of per-class recalls), which is not inflated by unequal class sizes, and was compared against a null distribution generated by 1,000 permutations of the class labels, each subjected to the full cross-validation procedure. Chance performance corresponds to 1/*k* for *k* classes.

### Categorical associations

The association between waveform class and RF class was assessed using Pearson’s chi-square test of independence. Expected frequencies were calculated from the marginal class totals, and degrees of freedom were defined as *df*=(*r*-1)(*c*-1). Effect size was quantified using Cramér’s V. Where expected cell counts fell below five, *p*-values were instead estimated by Monte Carlo permutation of the class labels (10,000 permutations).

### Controlling for recording penetration

Units recorded in the same penetration share eccentricity, electrode and stimulus, and are therefore not independent samples. To test each classification’s independent contribution within penetrations, we fitted a PERMANOVA in which penetration indicators and the other classification were entered as covariates before the class factor of interest. Class labels were permuted only among units of the same penetration (5,000 permutations), and effect sizes are reported as partial ω². To test whether class could be decoded in recordings the classifier had not seen, we repeated the linear discriminant classification with leave-one-penetration-out and leave-one-animal-out cross-validation, scoring balanced accuracy. Significance was assessed against null distributions generated by permuting class labels either across all units or within penetrations, each followed by the full cross-validation procedure (1,000 permutations).

## Results

### Extracellular spike waveform classes

A total of 226 RF-mapped single units were recorded from electrode penetrations in the LGN of three macaques (MCH, *n* = 88; MST, *n* = 26; MVG, *n* = 112). Mean extracellular spike waveforms were classified using a graph-based clustering of their principal components (see Methods), which yielded four distinct waveform classes (Figure 2A, B). Two classes exhibited typical negative-spiking waveforms (a dominant negative deflection followed by a smaller, slower positive deflection), which we defined as *Narrow* (Nar, 34%) and *Broad* (Bro, 21%) following the literature that negative-spiking waveforms separate into narrow-and broad-spiking populations commonly associated with inhibitory and excitatory neurons (Henze *et al*., 2000; Barthó *et al*., 2004). The two negative-spiking classes differed primarily in their trough-to-peak duration and the magnitude of their positive deflection: Broad waveforms had a much longer trough-to-peak duration (duration *ε*² = 0.22, *p* < 0.001; Figure 2C), and Narrow waveforms a larger trough-to-peak ratio, although the effect was small (TP-ratio *η*² = 0.05, *p* < 0.05).

**Figure 2.**
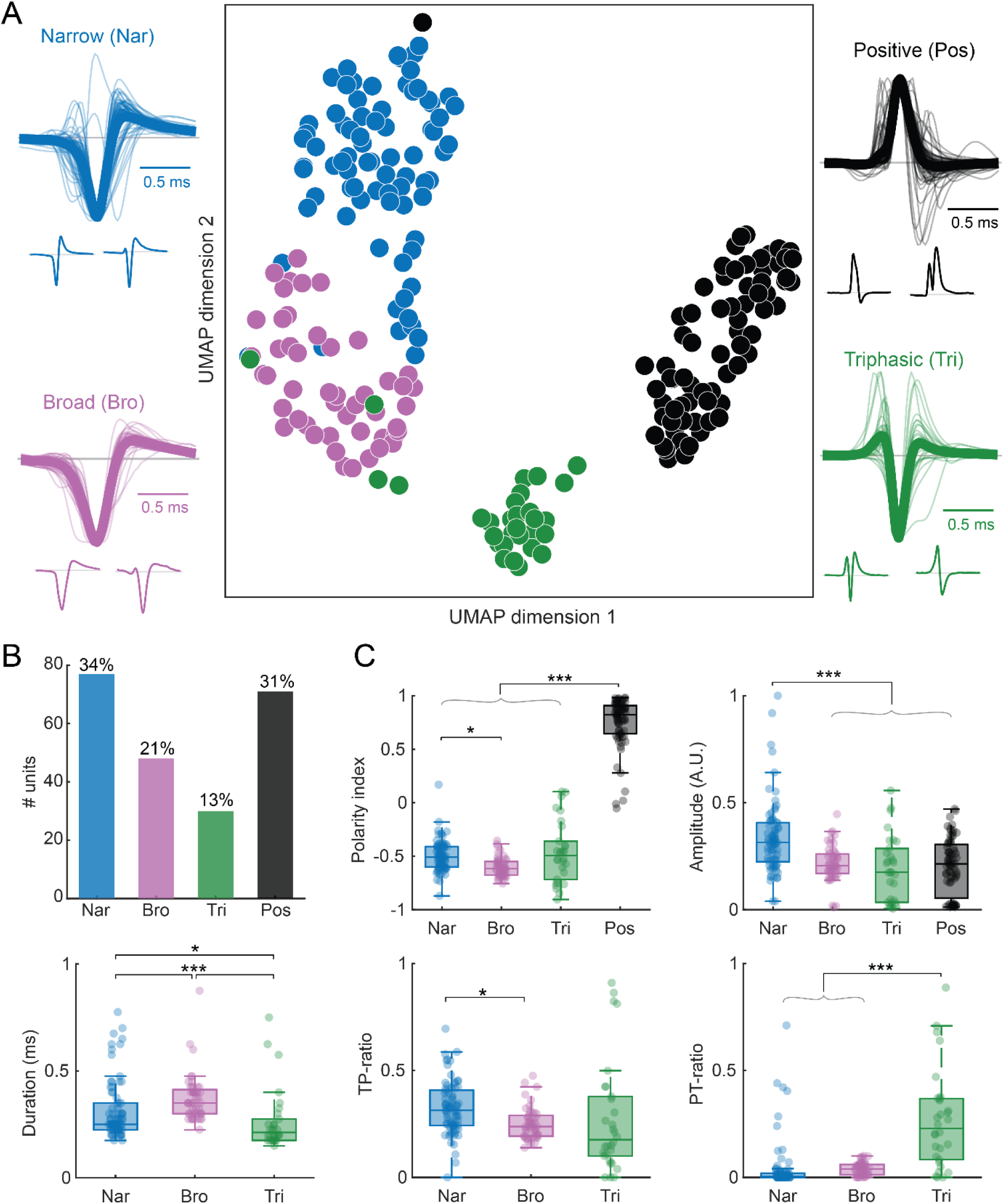
Graph-based classification of extracellular spike waveforms. **(A)** UMAP representation of mean extracellular spike waveforms from 226 RF-mapped LGN units, colored by waveform class: Narrow (Nar, blue), Broad (Bro, purple), Triphasic (Tri, green), and Positive (Pos, black). UMAP was calculated from the 20 waveform principal components used for graph construction and clustering. Surrounding plots show the processed mean waveform of each unit as a light trace and the class-average waveform as a thick trace. Two representative waveforms are shown beneath each class distribution. Waveforms were baseline-subtracted and normalized by their maximum absolute amplitude. Scale bars, 0.5 ms. **(B)** Number and percentage of units assigned to each waveform class. **(C)** Distributions of waveform polarity index, normalized amplitude, trough-to-post-peak duration, trough-to-post-peak ratio (TP-ratio), and pre-peak-to-trough ratio (PT-ratio). Duration, TP-ratio, and PT-ratio were not evaluated for positive-dominant waveforms. Boxes show the interquartile range, center lines show medians, and whiskers extend to the most extreme observations within 1.5 times the interquartile range. Points show individual units, including observations beyond the whiskers. Brackets identify significant FDR-corrected post hoc pairwise comparisons. \**p_FDR_*<0.05, \*\**p_FDR_*<0.01, \*\*\**p_FDR_*<0.001.

The third class, *Triphasic* (Tri, 13%), was also negative-spiking but carried an additional initial positive deflection preceding the primary negative deflection (Figure 2A), giving the shortest duration (*p* < 0.05) and the largest pre-peak-to-trough ratio of the negative-spiking classes (PT-ratio *η*² = 0.32, *p* < 0.001). The fourth class, *Positive* (Pos, 31%), was positive-spiking, with a dominant positive deflection that clearly separated it from the negative-spiking classes on the polarity index (*ε*² = 0.69, *p* < 0.001; Figure 2C). Narrow waveforms had the largest mean amplitude of the four classes (*η*² = 0.17, *p* < 0.001), suggesting that Narrow units may represent recordings from large neurons or cells near the electrode (Holt & Koch, 1999; Gold *et al*., 2006). Illustrations of all extracellular mean spike waveforms and the clustering are provided in Figures S3 and S4.

Positive and triphasic waveforms, although less commonly reported than negative waveforms, have been linked to activity distal to the soma: dendritic activity from return (capacitive) currents (Holt & Koch, 1999; Gold *et al*., 2006; Einevoll *et al*., 2013; Gulyás *et al*., 2016; Bakkum *et al*., 2019; Emmenegger *et al*., 2019; Bestel *et al*., 2021); action potentials propagating through axonal fibers <u>(Hubel, 1960; Bishop *et al*., 1962; Heinricher, 2004; Meeks *et al*., 2005; Lewandowska *et al*., 2015; Barry, 2015</u>; reviewed in <u>Sibille *et al*., 2026; Sun & Ibbotson, 2026)</u>; and afferent fibers originating from preceding processing areas (Robbins *et al*., 2013; Sun *et al*., 2021; Sibille *et al*., 2022, 2024), i.e., the retina in the present case. A recent high-density study of macaque V1 similarly resolved narrow-, broad-, and triphasic-spiking classes and found positive-spiking waveforms to be concentrated in the white matter, consistent with an axonal origin (Carr *et al*., 2026). We return to this axonal interpretation below, where the multivariate analyses provide convergent, but not definitive, evidence.

### RF classification

All 226 units were presented with chromatic noise stimuli to estimate their chromospatiotemporal receptive field (RF) by spike-triggered averaging (Figure 3A; see Methods). Fixation during mapping was stable, with gaze varying well within the fixation window (Figure S4). RFs were classified by their relative long- (L), medium- (M), and short-wavelength (S) cone weights into three RF classes, which we label by their functional cone signatures rather than by the anatomical magnocellular/parvocellular/koniocellular pathways, because we did not obtain histological confirmation of pathway identity. Rather than assigning classes from the sign of the cone weights alone, we required each weight to exceed the cone-weight noise before it counted toward a unit’s class. We adopted this criterion because we found that the classical sign-only method frequently misclassified RFs: several units that qualitatively resembled LM (magnocellular-type) were labeled LM-opposing (parvocellular-type) because weak L- or M-cone responses within the noise floor were treated as genuine opponency. Applying these noise thresholds, units whose S-cone weight was dominant or exceeded the cone-weight noise by more than three standard deviations were classified as S (10%; RF 4 in Figure 3A); units whose L- and M-cone weights were opposing and each exceeded the cone-weight noise by more than two standard deviations were classified as LM-opposing (LM-opp, 25%; RF 2 and 3); and the remaining units, with same-signed L- and M-cone weights, were classified as LM (65%; RF 1). We held S RFs to the stricter three-standard-deviation floor because a spurious S-cone signal is more likely to be mistaken as a distinct class than a weak L/M imbalance is. These noise-based thresholds brought the automated classification into closer agreement with qualitative inspection of the RF maps. This functional classification is closely related to the anatomical classifications of prior LGN studies (Wiesel & Hubel, 1966; Derrington & Lennie, 1984; White *et al*., 1998; Reid & Shapley, 2002).

**Figure 3.**
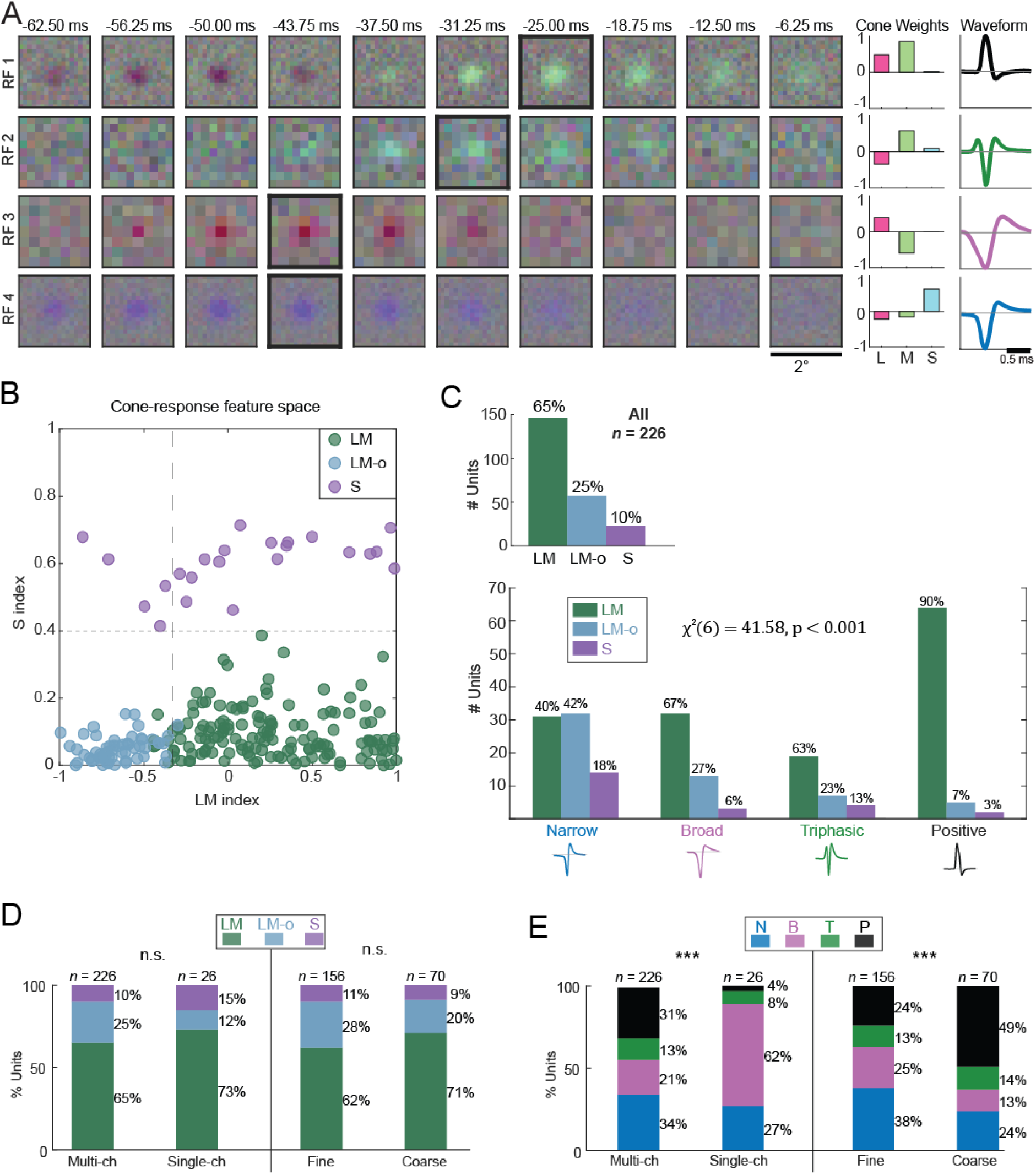
RF classification and its association with waveform class. **(A)** Four example chromospatiotemporal RFs shown across ten stimulus frames preceding the spike, from −62.50 to −6.25 ms in 6.25-ms increments. The thick black outline identifies the frame containing the maximum RGB RF-vector magnitude. Normalized L-, M-, and S-cone weights and the corresponding mean extracellular waveform are shown to the right of each RF. Scale bars, 2° and 0.5 ms. **(B)** RFs plotted in the continuous feature space defined by the LM index and S index and colored by RF class. LM RFs were characterized primarily by same-sign L-and M-cone contributions, LM-opposing RFs by significant opposite-sign L- and M-cone contributions, and S RFs by prominent S-cone contributions. Dashed lines are visual references and do not independently define the RF classes. **(C)** Top, number and percentage of the 226 units assigned to each RF class. Bottom, RF-class composition of each waveform class. Percentages indicate the proportion of each RF class within a waveform class. Waveform and RF class were significantly associated by Pearson’s chi-square test. **(D)** RF-class composition of units obtained using multichannel or single-channel electrodes and of multichannel units mapped using fine- or coarse-resolution stimuli. **(E)** Waveform-class composition of the same recording and stimulus subsets. Percentages indicate within-subset proportions, and *n*indicates the number of units in each subset. Population-composition p-values were obtained from contingency-table tests and corrected together using the Benjamini–Hochberg procedure. n.s., not significant; \*\**p_FDR_*<0.01; \*\*\**p_FDR_*<0.001.

The normalized cone weights of all 226 RFs occupied a cone-response feature space defined by an L/M-opponency index and an S index (Figure 3B). The distributions overlapped but showed structured separation, with S-cone units separated at an S-index threshold of 0.4 and LM-opponent units separated from LM units at an L/M-opponency index of −0.33, as indicated by the horizontal and vertical reference lines (descriptive visual guides only). The LM index showed a weak positive correlation with RF quality, measured by RF *z*-score (Pearson’s *r*=0.22, *p*<0.001), but was not correlated with waveform quality measured by SNR (*r*=-0.07, *p*=0.319). Thus, RF quality accounted for only a small proportion of the variability in the LM index. The S index was not significantly correlated with either RF *z*-score (*r*=0.09, *p*=0.193) or waveform SNR (*r*=0.10, *p*=0.117). S units predominantly displayed S-cone *on* responses, with roughly a 3:1 ratio of S-*on* to S-*off* RFs, consistent with prior work in marmoset LGN (Martin & Lee, 2014; Pietersen *et al*., 2014) and macaque retinal ganglion cells (De Monasterio *et al*., 1975; De Monasterio & Gouras, 1975).

Although absolute anatomical depth could not be compared reliably across sessions, relative depth was examined within nine approximately linear penetrations. In eight of nine penetrations (89%), the mean relative depth of LM units was at or below the mean depths of both LM-opposing and S units. This within-penetration pattern was descriptive and was not subjected to a pooled depth analysis because recording trajectories, entry positions, and anatomical layer boundaries could not be reconstructed consistently across sessions.

### Associations between waveform and RF classes

Waveform and RF class were significantly associated [*χ*²(6, *N* = 226) = 41.58, Monte Carlo *p* < 0.001, Cramér’s *V*=0.30; Figure 3C]. The association was driven by several converging patterns: the positive-spiking class was overwhelmingly LM (Pos: 90% LM, 7% LM-opposing, 3% S), whereas the negative-spiking classes carried a larger share of LM-opposing and S RFs (Nar: 40% LM, 42% LM-opposing, 18% S; Bro: 67% LM, 27% LM-opposing, 6% S; Tri: 63% LM, 23% LM-opposing, 13% S). This heterogeneity may reflect genuine functional differences between waveform classes, but may also reflect sampling biases inherent to extracellular recording (Towe & Harding, 1970) and anatomical layout (see Discussion).

To assess whether recording- or stimulus-related sampling biases contributed to this heterogeneity, we compared class composition across recording and stimulus subsets. RF-class composition did not differ significantly between multichannel and single-channel recordings or between fine- and coarse-resolution stimuli (both n.s.; Figure 3D), whereas waveform-class composition differed across both comparisons (both *p* < 0.001, FDR-corrected; Figure 3E). Waveform-class sampling was thus more sensitive than RF-class sampling to electrode configuration and stimulus resolution. In particular, the high proportion of LM RFs among positive-spiking units is compatible with preferential sampling of non-somatic signals, including retinal afferents traversing or terminating within LM-dominated magnocellular layers, but the recorded compartment could not be identified from these data.

### Waveform and RF classes occupy a shared physiological feature space

The categorical association above shows that waveform and RF labels are related, but not whether they capture the same underlying structure or independent aspects of LGN organization. To address this, we analyzed both classifications within a physiological feature space comprising RF eccentricity, RF latency, spike rate, burst index, CV_2_ of the inter-spike interval, ACG peak latency, ACG refractory recovery, and RF *z*-score (156 units with complete measurements on all features; Nar *n* = 60, Bro *n* = 39, Tri *n* = 20, Pos *n* = 37; LM *n* = 96, LM-opposing *n* = 43, S *n* = 17). The L/M and S indices were excluded to avoid circularity, as the RF classes are defined by them.

Projected onto the first two principal components (PCs), both waveform classes and RF classes occupied overlapping but non-identical regions of the feature space (Figure 4 A,B; PC1 and PC2 together captured 55.5% of the variance, so the two-dimensional projection understates the separation quantified below). Permutational multivariate analysis of variance confirmed that class membership accounted for a significant fraction of the multivariate variation: waveform class explained 19.5% of the variance (PERMANOVA, *R*² = 0.195, *F*(3, 152) = 12.3, *p* = 0.0001) and RF class 11.0% (*R*² = 0.110, *F*(2, 153) = 9.4, *p* = 0.0001). Because PERMANOVA is sensitive to differences in multivariate dispersion as well as location, we tested homogeneity of dispersions (PERMDISP): dispersions were homogeneous across RF classes (*p* = 0.52), so that result reflects a difference in centroid location, whereas dispersions differed across waveform classes (*p* < 0.001; Figure S5), the Broad class forming a more compact cluster than the others. We therefore interpret the waveform result alongside the classification analyses below, which do not depend on dispersion.

**Figure 4.**
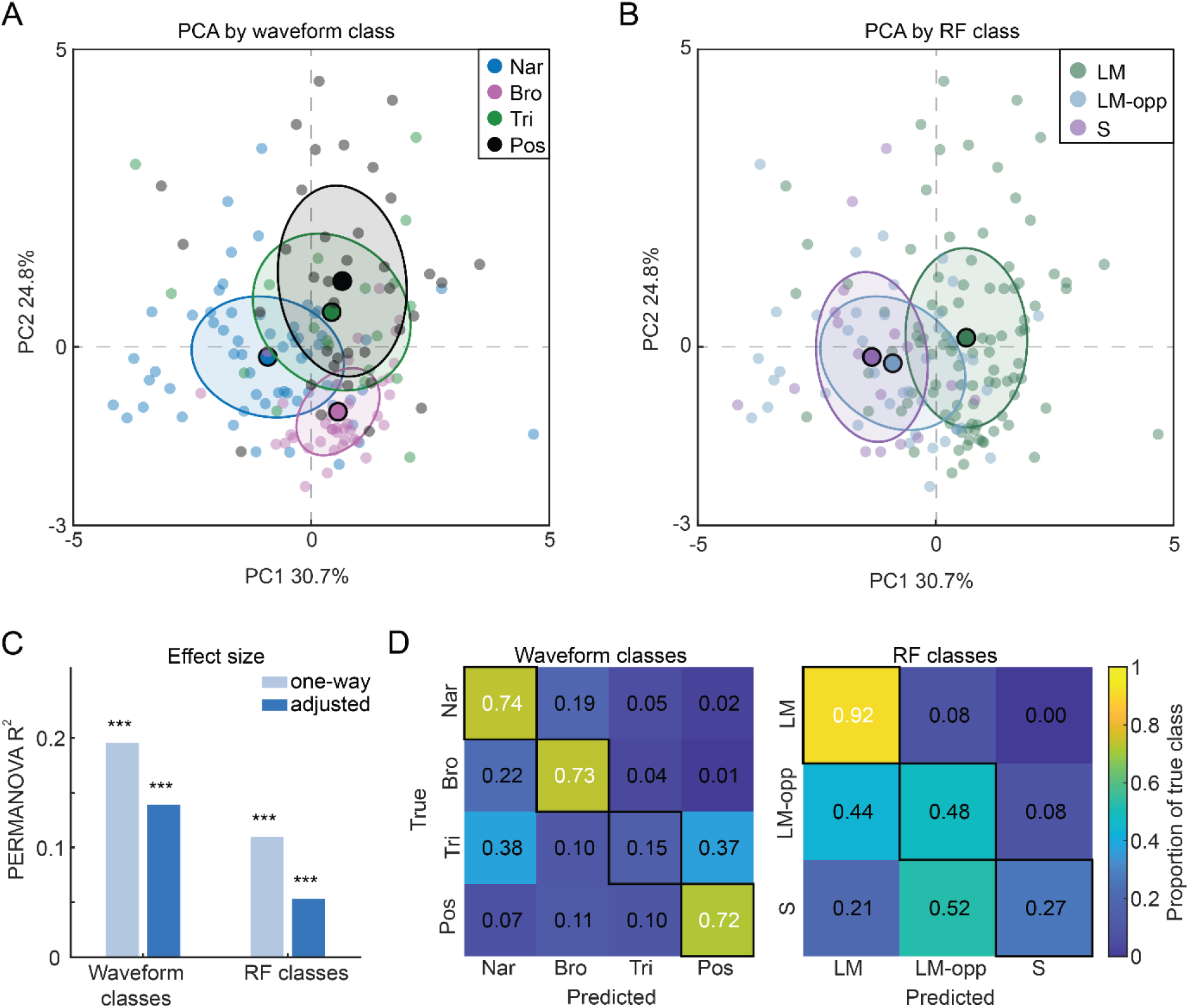
Waveform and RF classes occupy partially independent regions of physiological feature space. **(A, B)** Principal-component representation of the shared physiological feature matrix, colored by waveform class (**A**) or RF class (**B**). The matrix contained RF eccentricity, RF latency, spike rate, burst index, CV_2_, ACG peak latency, ACG refractory recovery, and RF z-score. Analyses included 156 units with complete measurements for all features: Nar, *n*=60; Bro, *n*=39; Tri, *n*=20; Pos, *n*=37; LM, *n*=96; LM-opposing, *n*=43; and S, *n*=17. Small points represent individual units, large outlined points indicate class centroids, and ellipses extend one standard deviation along the principal covariance axes of each class. Axis labels report the percentage of total variance explained by each principal component. **(C)** PERMANOVA effect sizes for waveform and RF classes. Light bars show one-way *R*^2^ values obtained when each classification was tested separately. Dark bars show partial *R*^2^ values from the two-factor model, representing the variance explained by each classification after accounting for the other. Significance was estimated using 10,000 permutations. **(D)** Row-normalized confusion matrices from regularized linear discriminant classifiers trained on the same standardized physiological features. Classification used stratified five-fold cross-validation repeated 20 times. Rows indicate true classes, columns indicate predicted classes, and cell values indicate the proportion of each true class assigned to each predicted class. Outlined diagonal cells represent per-class recall. \*\*\**p_perm_*<0.001.

Two features of these analyses indicate that the waveform and RF schemes capture partially independent structure. First, in a two-way model that fitted both factors jointly, waveform class explained variance beyond that accounted for by RF label (partial *R*² = 0.139, *F*(3, 150) = 9.3, *p* < 0.0001), and RF label explained variance beyond waveform class (partial *R*² = 0.053, *F*(2, 150) = 5.3, *p* < 0.0001). Second, a cross-validated classifier trained on the same features recovered waveform class at 58% balanced accuracy (chance 25%, permutation null mean 0.25; *p* < 0.001) and RF class at 55% (chance 33%, null mean 0.33; *p* < 0.001), confirming that class membership is predictable from the physiological features and not merely a difference in spread.

Neither classification was equally separable across all classes, and the pattern of separability is itself informative (Figure 4). Among waveform classes, all pairs were distinguishable except Triphasic and Positive (pairwise PERMANOVA, all *p_FDR_* < 0.05 except Tri-Pos, *R*² = 0.03, *p* = 0.15; Figure S5): the classifier recovered Narrow, Broad, and Positive units well (per-class recall 0.74, 0.73, and 0.72, respectively) but failed to recover Triphasic units (recall 0.15), which were assigned instead to Narrow (0.38) and Positive (0.37). The physiological indistinguishability of triphasic and positive waveforms, together with the leakage of triphasic units toward the positive class, is consistent with a shared non-somatic basis for these two non-classical waveform types <u>(Sibille *et al*., 2024; Carr *et al*., 2026; reviewed in Sun & Ibbotson, 2026)</u>. Among RF classes, LM units separated from both LM-opposing and S units (both pairwise *p* < 0.05), but LM-opposing and S units were not distinguishable from one another (*R*² = 0.02, *p* = 0.24; Figure S5): the classifier recovered LM units almost perfectly (recall 0.92) but performed poorly on LM-opposing (0.48, with 0.44 misassigned to LM) and S units (0.27, with 0.52 misassigned to LM-opposing). Thus the RF separation in feature space is carried predominantly by LM versus non-LM units, rather than by three fully distinct physiological classes.

Because RF eccentricity partly reflects electrode placement rather than intrinsic physiology, we repeated all multivariate analyses with eccentricity excluded. The waveform result was reduced but preserved (PERMANOVA *R*² = 0.148; cross-validated accuracy 0.52 versus 0.25 chance, *p* < 0.001; waveform-beyond-cone partial *R*² = 0.096, *p* < 0.0001), and the same two non-separable pairs (Triphasic–Positive and LM-opposing–S) were obtained, indicating that the separation is not carried by eccentricity alone.

As units from the same penetration can share eccentricity, electrode and stimulus, we repeated the analyses while controlling for recording penetration. With penetration controlled (penetrations entered as a blocking term and permutations restricted within penetrations; see Methods), waveform class still explained structure beyond RF class (partial ω² = 0.037, p < 0.001), and RF class beyond waveform class (partial ω² = 0.038, p < 0.001). Classifiers tested on penetrations or animals withheld from training remained above chance for both waveform class (balanced accuracy 0.59 and 0.48; chance 0.25) and RF class (0.56 and 0.46; chance 0.33; all p ≤ 0.005).

### Feature-level differences between classes

To identify which physiological features underlie the multivariate separation, we compared each feature across waveform and RF classes (Figure 5A, B; Benjamini–Hochberg–corrected across the feature family). Across waveform classes, differences were widespread: RF eccentricity (*ε*² = 0.47), RF latency (*η*² = 0.34), ACG refractory recovery (*ε*² = 0.27), ACG peak latency (*ε*² = 0.19), burst index (*ε*² = 0.18), CV_2_ (*η*² = 0.16), spike rate (*ε*² = 0.09), and RF *z*-score (*η*² = 0.05) all differed significantly (all *p_FDR_*<0.05). Positive and Triphasic units had RFs at markedly larger eccentricities (median 16.5° and 11.3°, respectively) than the negative-spiking Narrow and Broad units (3.6° and 3.0°), and RF latency decreased across the classes from Narrow (median 43.8 ms) to Positive (25.0 ms). Part of this difference reflected RF class and stimulus: restricted to LM units recorded with the same stimulus (β = 1.5), Positive units still responded earlier than Narrow units (median 31.25 vs 37.5 ms, n = 13 and 19, p < 0.001). Among the negative-spiking classes, Narrow units had the lowest burst index and the longest autocorrelogram peak latency and refractory recovery, whereas Broad units fired at the lowest rate and most irregularly (highest CV_2_; Figure 5A). Across RF classes, RF latency (*η*² = 0.34), burst index (*ε*² = 0.23), ACG refractory recovery (*ε*² = 0.23), ACG peak latency (*ε*² = 0.09), and eccentricity (*ε*² = 0.07) differed significantly, while spike rate, CV_2_, and RF *z*-score did not (*p_FDR_*>0.05); LM units had shorter RF latencies (median 31.3 ms, versus 43.8 ms and 50.0 ms for LM-opposing and S units), higher burst indices, shorter autocorrelogram peak latencies, and faster refractory recovery than LM-opposing and S units, consistent with the known temporal properties of the magnocellular-associated pathway (Maunsell *et al*., 1999; Pietersen *et al*., 2014; Eiber *et al*., 2018).

**Figure 5.**
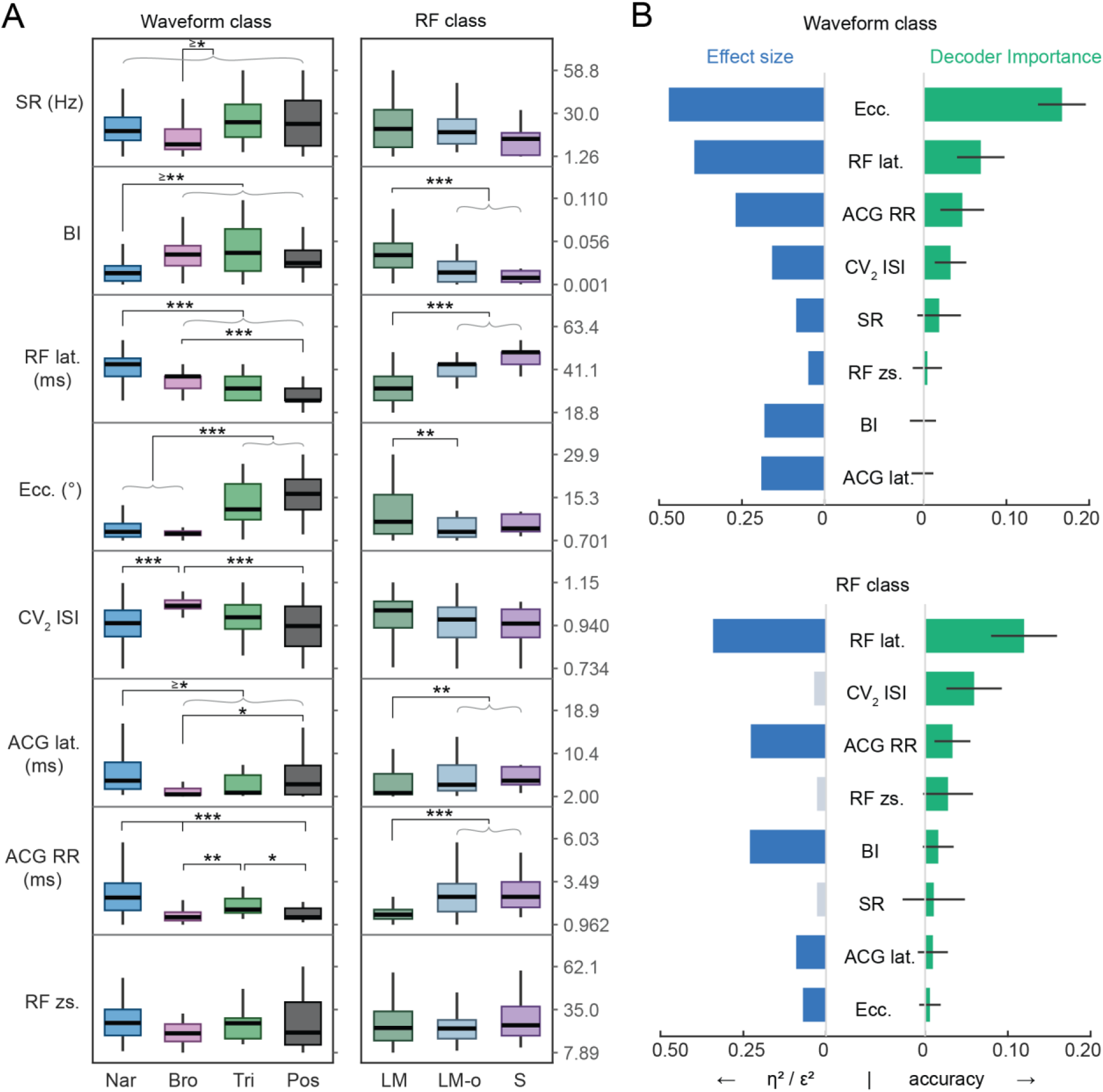
Physiological features associated with waveform and RF classes. **(A)** Compact boxplot matrices showing spike rate (SR), burst index (BI), RF latency, RF eccentricity, local interspike-interval variability (CV_2_ ISI), ACG peak latency, ACG refractory recovery (ACG RR), and RF z-score across waveform classes and RF classes. All finite observations meeting the inclusion criteria were retained; sample size may therefore differ among metrics according to data availability. Within each row, values were mapped from the pooled 2nd to 98th percentile to permit metrics with different units and ranges to be displayed together. The three numerical references at the right of each row indicate the lower endpoint, midpoint, and upper endpoint of this display range. Boxes show the interquartile range, center lines show medians, and whiskers extend to the most extreme observations within 1.5 times the interquartile range. Brackets identify significant post hoc comparisons after FDR correction within each metric. **(B)** Comparison of univariate effect size and multivariate decoder importance. Blue bars extending left show *η*^2^ for ANOVA or Welch ANOVA and *ε*^2^ for Kruskal-Wallis tests. Dark blue indicates a significant FDR-corrected omnibus test, whereas pale blue indicates a nonsignificant test. Green bars extending right show permutation feature importance, defined as the reduction in held-out balanced accuracy after shuffling that feature while retaining the remaining features. Gray bars indicate importance estimates at or below zero. Error bars show the standard deviation across 20 repetitions of five-fold cross-validation. Metrics are ordered separately for waveform and RF classification according to decoder importance.

Because the univariate tests treat each feature in isolation, we additionally quantified how much each feature contributed to class separation once the others were accounted for, using permutation feature importance from the cross-validated classifier (Figure 5B). For both classifications, the univariate effect sizes and the multivariate importance measures diverged in informative ways. Burst index differed strongly across classes on its own (*ε*^2^=0.18 for waveform and *ε*^2^=0.23 for RF) yet contributed almost nothing to decoding once the other features were included (importance = −0.01 and 0.02, respectively), indicating that its class differences are largely redundant with the autocorrelogram-based timing measures. Conversely, CV_2_ did not differ significantly across RF classes on its own (*η*² = 0.04, n.s.) but was the second most important feature for decoding RF class (importance = 0.07, versus 0.13 for RF latency), indicating that it carries information about class only in combination with the other features. RF eccentricity was the single strongest contributor to waveform-class separation by both measures (*ε*^2^=0.47² = 0.47; importance = 0.17, more than twice the next feature), consistent with its role in the sensitivity analysis above. These divergences show that univariate significance and multivariate contribution are distinct, and that no single feature accounts for the class structure; rather, LGN organization along both waveform and RF labels is distributed across multiple extracellular and functional measurements.

## Discussion

In this study, we surveyed extracellular spike waveform diversity among RF-mapped units in the LGN of awake rhesus macaques and examined whether waveform-defined classes were associated with RF and spiking properties. Graph-based clustering of waveform shape alone identified four graph-derived classes (Narrow, Broad, Triphasic, and Positive) that subsequently differed in RF class composition, response timing, firing statistics, and autocorrelation properties. Waveform and RF classes were significantly associated, yet each explained independent variation in the shared physiological feature space. Extracellular waveform shape and RF response profile therefore describe related but partially distinct aspects of LGN organization, and neither resolves the population into discrete, one-to-one cell types.

The classes were not fully separable. Narrow, Broad, and Positive units were recovered reasonably well from their physiological features, whereas Triphasic units overlapped with both Narrow and Positive units, and LM units were more readily distinguished than LM-opposing and S units. We therefore interpret the classes as structured, overlapping regions of a continuous feature space rather than discrete biological types, consistent with recent work showing that electrophysiological identities often form continua and are better described by combinations of waveform, firing, and temporal properties than by any single measurement (Lee *et al*., 2026).

### Extracellular waveform diversity

The Narrow and Broad classes carried the negative-dominant waveform traditionally associated with recordings near the soma and axon initial segment; they differed mainly in trough-to-post-peak duration, amplitude, and post-trough positivity, variation of the kind expected from differences in ion-channel kinetics, cellular morphology, and the position of the recording contact, which recent reviews describe in detail (Sibille *et al*., 2026; Sun & Ibbotson, 2026; Siegle & Steinmetz, 2026). We did not interpret these classes simply as inhibitory and excitatory neurons. Although the distinction is frequently used, narrow spikes can arise from excitatory neurons in macaque cortex and some inhibitory neurons can produce broad spikes (Vigneswaran *et al*., 2011; Lemon *et al*., 2021; Shaw *et al*., 2026), and recent Neuropixels recordings in macaque V1 recovered more narrow-spiking units than the parvalbumin-positive population can account for (Carr *et al*., 2026). Waveform width is therefore physiologically informative but insufficient for assigning excitatory or inhibitory identity.

The Triphasic and Positive classes are less compatible with a simple somatic source. Both are dominated by, or begin with, a positive deflection, which may reflect dendritic return currents or the current sources flanking an action potential as it propagates along an axon (Sibille *et al*., 2026; Sun & Ibbotson, 2026). Several observations are consistent with a non-somatic, potentially axonal, origin: both classes had the shortest RF latencies and the largest eccentricities; Positive units were strongly enriched for LM RFs (90%); and the two classes were not significantly separated in the pairwise multivariate comparison, with Triphasic units frequently predicted as Positive by the classifier. This is the pattern expected if both reflect fibers whose recorded polarity varies with electrode location relative to local current sources and sinks. Positive and triphasic waveforms have been associated with afferent fibers and other non-somatic sources across preparations (Robbins *et al*., 2013; Sun *et al*., 2021; Sibille *et al*., 2022, 2024), and positive waveforms concentrate in white matter in macaque V1 while negative waveforms predominate in gray matter (Carr *et al*., 2026). If positive units were fibers sampled at random, their RF classes would mirror retinal composition, which is dominated by midget cells; instead, 90% were LM. We therefore propose that many Positive units are parasol-cell axons, whose large diameter would make them preferentially detectable as they course through the LGN. We nonetheless regard these as candidate non-somatic populations rather than confirmed retinal afferents, because a single-channel waveform representation cannot separate relay-cell axons, dendritic currents, and mixed compartmental signals, or uniquely identify the contributing compartment.

### RF classes and their relationship to waveform classes

RFs were classified as LM, LM-opposing, or S according to their cone-response structure. We used these descriptive labels rather than magnocellular (M), parvocellular (P), and koniocellular (K) because anatomical pathway identity was not confirmed: cone-response profile and anatomical pathway are not equivalent, M and P populations overlap in several response properties, and K neurons are particularly heterogeneous (Wiesel & Hubel, 1966; Derrington & Lennie, 1984; White *et al*., 1998; Hendry & Reid, 2000; Reid & Shapley, 2002; Tailby *et al*., 2008). The continuous distribution of units in LM- and S-index space is consistent with our RF nomenclature: the three classes summarize response structure without forming perfectly discrete populations. The physiological comparisons nonetheless recovered expected differences. LM units had shorter RF latencies, higher burst indices, shorter ACG peak latencies, and faster refractory recovery than LM-opposing and S units — properties commonly associated with the magnocellular pathway (Maunsell *et al*., 1999; Pietersen *et al*., 2014; Eiber *et al*., 2018) — and RF latency was the strongest multivariate predictor of RF class. LM-opposing and S units, by contrast, overlapped substantially in both pairwise analyses and classification, and an exploratory within-penetration analysis placed LM units at or below the other classes in eight of nine approximately linear penetrations, broadly compatible with the ventral position of the magnocellular layers. The separation we detect is therefore carried mainly by an LM-versus-rest distinction; absolute depth and layer boundaries could not be reconstructed reliably, so we treated this pattern descriptively and did not use it to assign M, P, or K identity.

Waveform and RF classes were strongly associated: Positive units were predominantly LM, whereas Narrow units contained the largest proportions of LM-opposing and S RFs, with Broad and Triphasic units were intermediate. The informative result is that this association was incomplete. Waveform class explained approximately 19% of the one-way multivariate variation and RF class approximately 11%; after accounting for their association in the joint model, waveform class retained a partial effect of approximately 14% and RF class approximately 5%. Because clustering used waveform shape alone and the LM and S indices were excluded from the shared analysis, this independence cannot be an artifact of testing groups on the features that defined them. Waveform morphology therefore captured physiological structure not explained by RF class, and the reverse also held; the two schemes intersect rather than coincide.

Some caution is warranted because multivariate dispersion differed among waveform classes: Broad units occupied a more compact region of feature space than the others (PERMDISP, *p* < 0.001), so both centroid location and within-class variation may have contributed. Waveform labels were nonetheless decoded above chance and most class pairs were significantly separated, indicating that the observed structure was not limited to the omnibus PERMANOVA result. No single measurement produced the structure. RF eccentricity and latency produced the largest univariate effects for waveform class, and because eccentricity partly reflects the locations sampled by the electrode, we repeated the multivariate analysis without it; separation and decoding remained significant, indicating the result was not solely a retinotopic sampling effect. Univariate effect size and multivariate importance did not always agree — burst index differed strongly among classes but added little unique decoding information, probably because it was correlated with ACG timing, whereas CV_2_ contributed to RF-class prediction despite a small univariate effect. Class identity was thus distributed across correlated extracellular and functional features, which is why an integrated treatment recovers it where any single feature cannot (Lee *et al*., 2026).

## Implications and limitations

These results bear on how the early visual pathway is characterized from extracellular recordings. Population descriptions of the LGN, including its chromatic tuning, response latencies, and the relative weights of the streams feeding V1, are assembled by counting units and reading off their properties, on the inferred assumption that each unit is a relay-cell soma.

Single-electrode recording has long been known to favor large, active, visually responsive somata and to discard the rest, so that the characterized population is a biased sample of the real one (Olshausen & Field, 2005). Our data add a second bias, orthogonal to the first and found in the recorded signal rather than the cell: if a substantial fraction of RF-mapped LGN units are axonal or otherwise non-somatic, extracellular surveys not only oversample certain somata but may fold afferent fibers into the same classifications, and do so unevenly. A survey that treats every unit as a relay soma would therefore bias estimates of both the composition and the timing of the geniculate signal arriving at V1. Waveform shape offers a simple, per-unit assessment for this concern, allowing the identification of units that warrant additional examination before being included in a population summary.

Consistent with these observations on recording bias, waveform-class composition varied with electrode type and stimulus resolution while RF-class composition did not, marking extracellular waveform sampling as the more bias-prone of the two measurements. Extracellular potentials decay rapidly with distance, and their shape depends on neuronal geometry, electrode properties, tissue motion, filtering, and contact position relative to neuronal compartments (Siegle & Steinmetz, 2026); dense arrays improve sampling but do not eliminate these biases, however, they also may reveal axonal or dendritic signals rarely detected with traditional electrodes.

We acknowledge that several experimental limitations constrain these conclusions. One, the session-specific fixation windows were large relative to some LGN RFs, which is why we excluded RF size from analysis (see Figure S4). Two, absolute depth and layer boundaries could not be compared consistently across sessions and were therefore used only descriptively. Three, the present results should be viewed as a survey of RF-mapped units meeting SNR and completeness criteria rather than a census of LGN cell types, and the smaller Triphasic and S groups were correspondingly less precisely characterized. Most fundamentally, these classes are electrophysiological populations rather than anatomically validated cell types: without histology, projection identification, or genetic or optogenetic ground truth, no waveform class can be bound to an anatomical cell type. High-density recordings that contain multichannel waveform for a given action potential source, and thus more completely measure each spike’s spatial footprint and propagation, are the clearest route to separating somatic, axonal, and dendritic sources rather than merely identifying potential candidates (Sibille *et al*., 2026; Carr *et al*., 2026).

In conclusion, extracellular spike waveforms in the awake macaque LGN formed four reproducible classes associated with RF structure and spiking physiology but not reducible to it. Waveform shape and RF response profile each captured structure the other missed, and no single feature accounted for either. Rather than marking discrete cell types, waveform diversity carries probabilistic information about how the LGN is organized, and is most useful when considered alongside RF, response-timing, firing, and autocorrelation measurements rather than on its own. More broadly, our findings underscore the importance of considering recording biases in extracellular studies and offer a practical framework for describing neuronal populations in primate and deep-brain recordings, where direct cell-type identification remains difficult.

## Acknowledgments

SHS, NJK, and JSP were supported by William M. Wood Foundation, the National Eye Institute award EY027888, the NIMA Foundation, and generous donations from Peter and Yayi Pezaris, Michael Gersh, and Don Good. JIL was supported by the VA Biomedical Laboratory Research and Development award BX005959, Novo Nordisk award 0065289, and the National Institute for Neurological Disease and Stroke award NS110575.

## Data availability statement

The data that support the findings of this study are available from the corresponding author upon reasonable request.

## Supplementary Figure Legends

**Figure S1.**
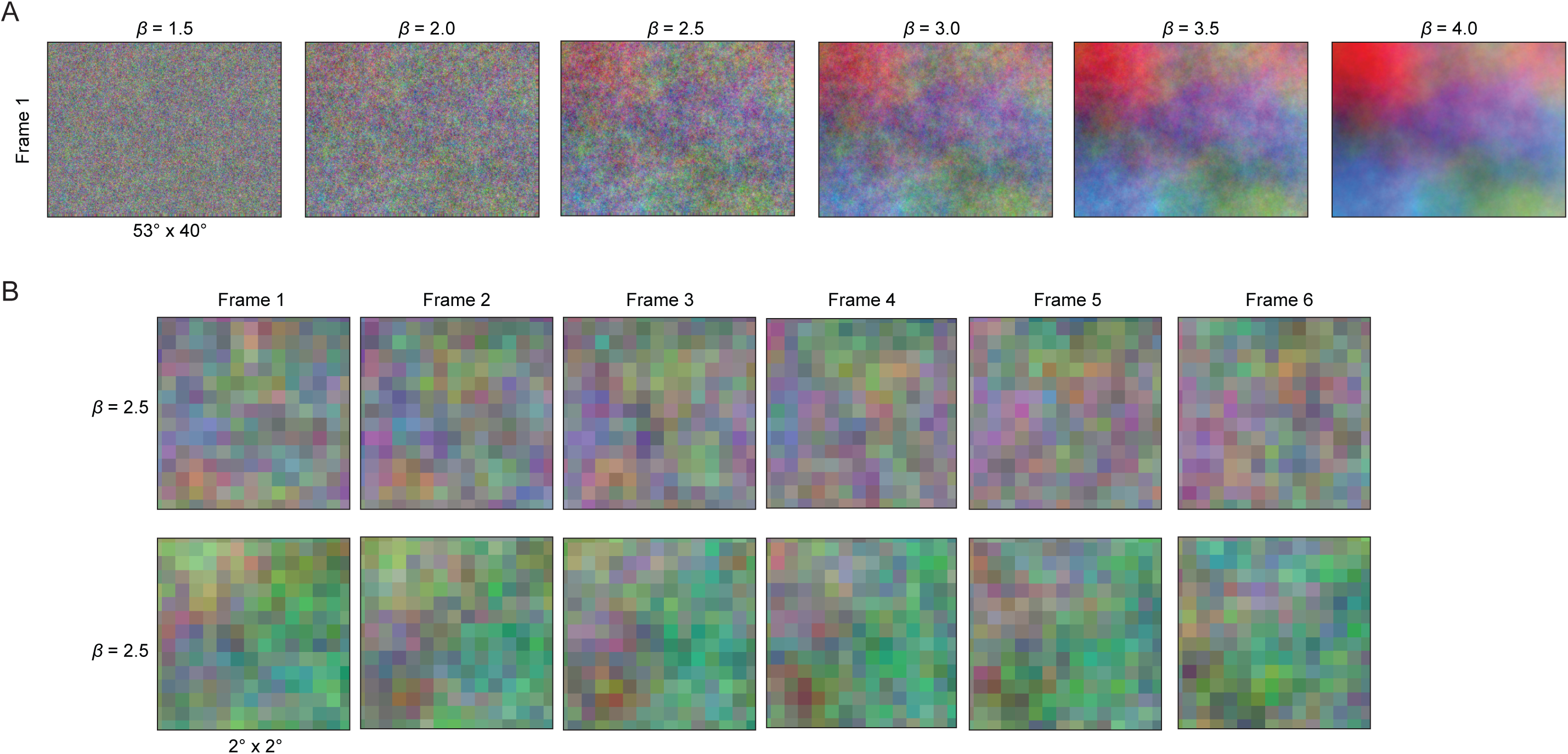
Colored noise stimuli used for RF mapping**. (A)** Example full-field colored noise frames generated with spatial power spectra proportional to 1/*f^β^*, at the indicated values of *β*. Increasing *β*produced stronger spatial correlations and greater relative power at low spatial frequencies. The full stimulus subtended approximately 53° × 40° of visual angle. **(B)** Six consecutive frames from example low-resolution (top) and high-resolution (bottom) colored noise sequences, displayed within 2° × 2° regions to illustrate their spatial and temporal structure.

**Figure S2.**
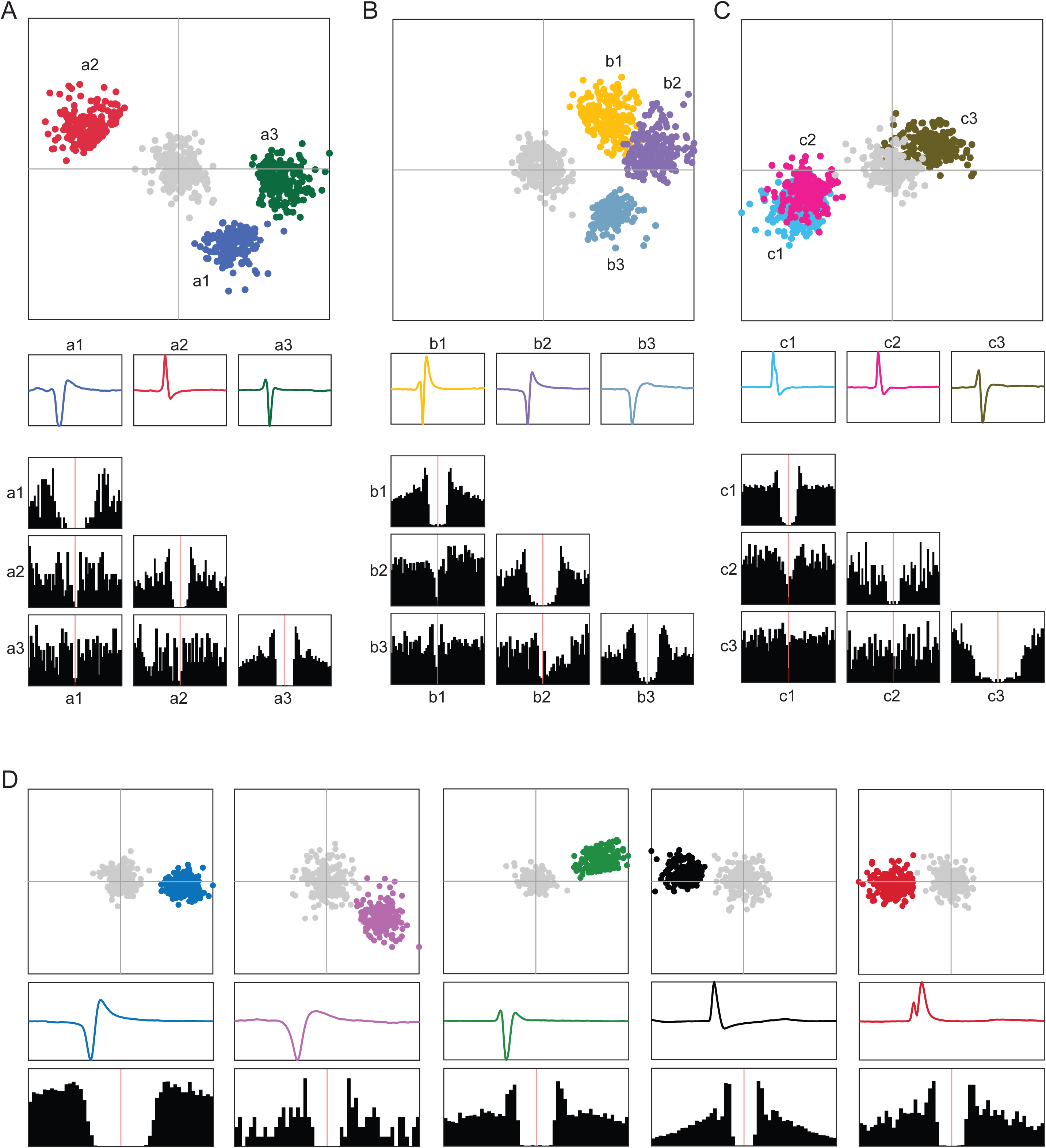
Representative spike-sorting examples from LGN recordings. **(A–C)** Three examples of simultaneously recorded spike clusters during manual curation. Top, clusters represented in principal-component space; colored points identify the highlighted clusters and gray points represent other detected events. Middle, mean extracellular spike waveform for each highlighted cluster. Bottom, autocorrelograms along the diagonal and pairwise cross-correlograms below the diagonal. Red vertical lines indicate zero time lag. Principal-component separation, waveform similarity, refractory structure, and cross-correlograms were considered together when determining whether clusters represented distinct single units or should be merged. **(D)** Five representative accepted single units spanning the waveform morphologies observed in the dataset. For each unit, the principal-component distribution relative to neighboring events, mean extracellular waveform, and autocorrelogram are shown from top to bottom. Colored points and traces identify the selected unit, whereas gray points indicate neighboring events.

**Figure S3.**
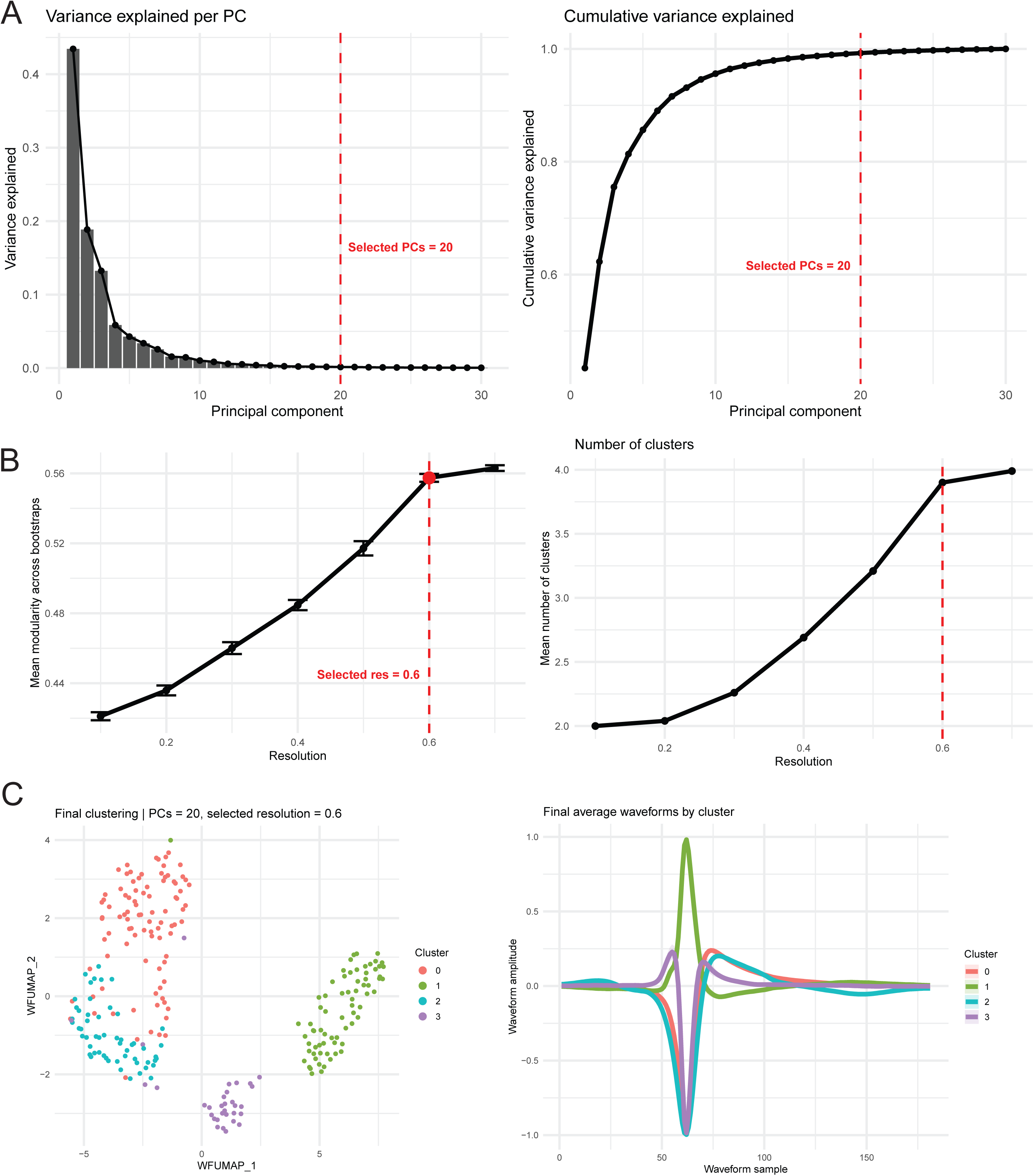
Selection of waveform principal components and clustering resolution. **(A)** Variance explained by each waveform principal component and cumulative variance explained across components. Red dashed lines indicate the 20 principal components selected for graph construction and clustering. **(B)** Mean graph modularity and mean number of recovered clusters across 100 bootstrap iterations at each tested clustering resolution. Each bootstrap used 80% of units sampled without replacement. Points and error bars summarize the bootstrap distributions. The selected resolution of 0.6 is indicated in red. **(C)** Final UMAP representation and average waveform associated with each graph-derived cluster using 20 principal components and a clustering resolution of 0.6. Graph-derived clusters were subsequently assigned the Narrow, Broad, Triphasic, and Positive waveform labels.

**Figure S4.**
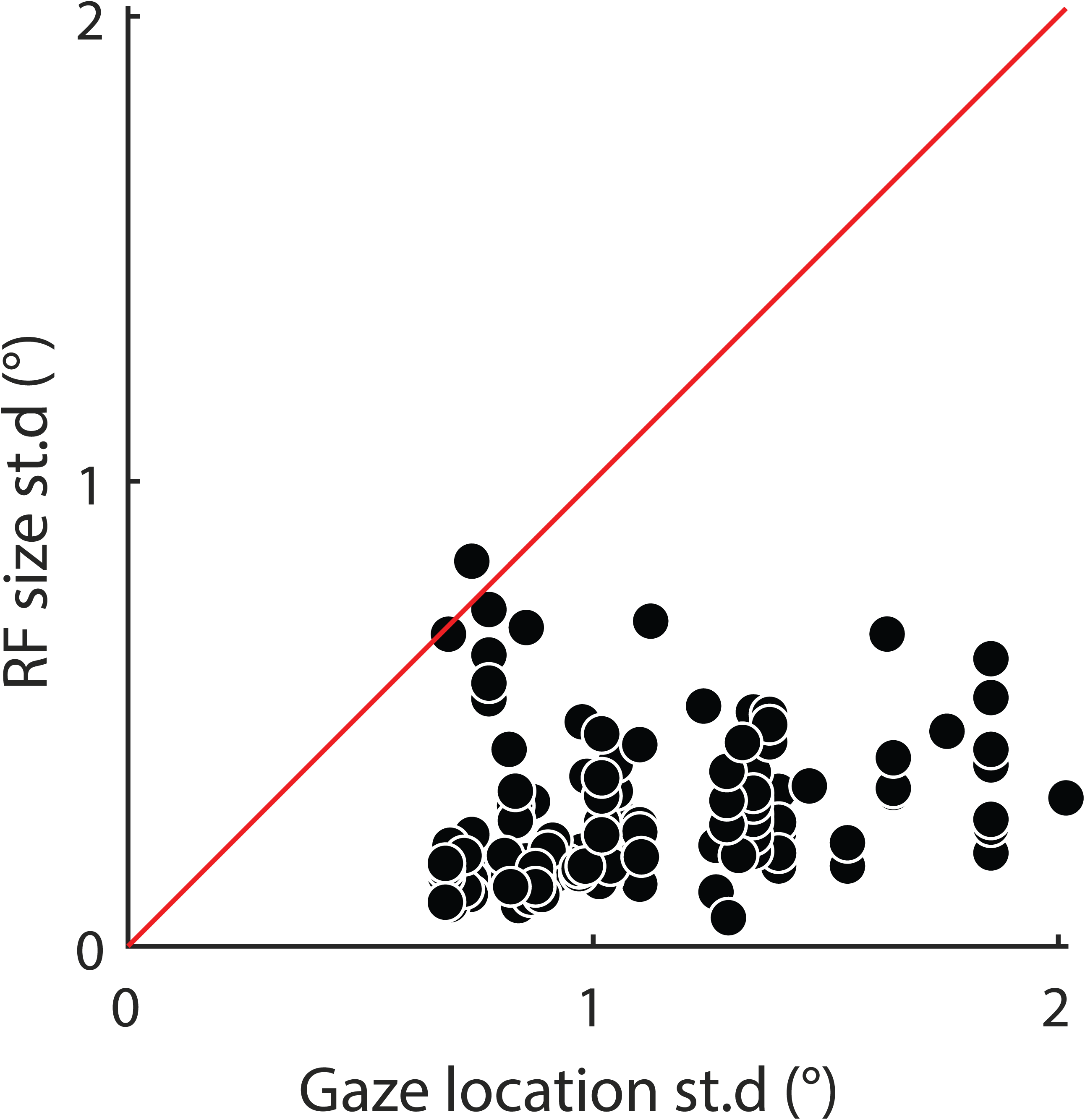
Relationship between gaze variability and estimated RF size. Standard deviation of estimated RF size plotted against the standard deviation of gaze position during RF mapping. Each point represents one RF for which both measurements were available. The red identity line represents equal RF-size and gaze-position variability, illustrating that typical fixation performance was substantially tighter than the 2° fixation window. With perfect, noiseless gaze position measurement, fixational eye movements would broaden the measured RFs, placing a lower bound on their apparent size. This plot shows that uncertainty in gaze position is substantially larger than RF size, suggesting that instrumentation noise dominates the measured variability in gaze position during fixation.

**Figure S5.**
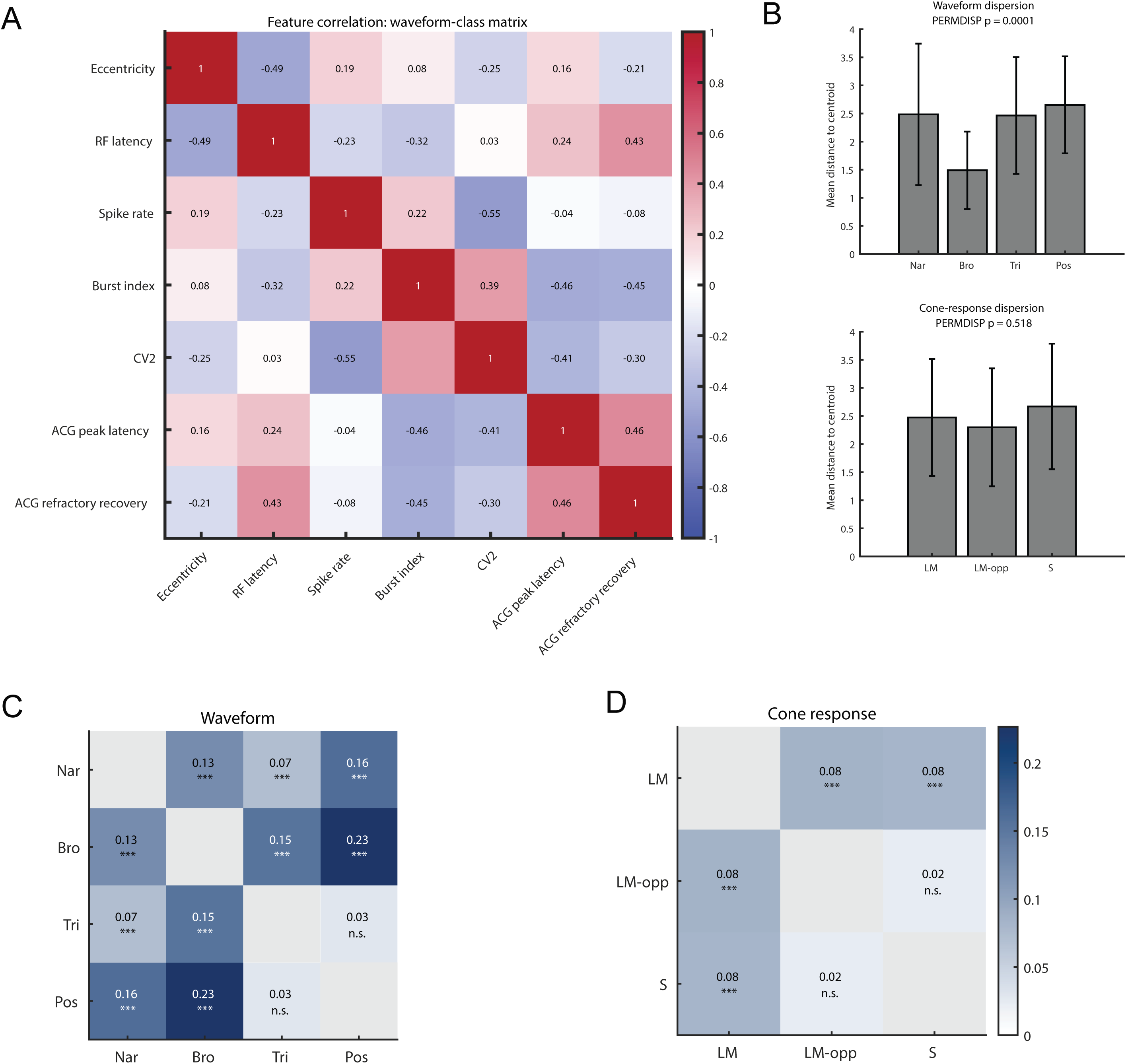
Feature correlations, multivariate dispersion, and pairwise class separation. **(A)** Pearson correlation matrix for RF eccentricity, RF latency, spike rate, burst index, CV_2_, ACG peak latency, and ACG refractory recovery. Values inside cells give Pearson correlation coefficients; warmer colors indicate positive correlations and cooler colors indicate negative correlations. RF z-score, which was included in the primary multivariate analysis, is not displayed in this panel. **(B)** Mean Euclidean distance to the class centroid for waveform classes and RF classes. Bars show group means, and error bars show standard deviations among units. PERMDISP tests used 10,000 permutations. **(C, D)** Pairwise PERMANOVA effect-size matrices for waveform classes (**C**) and RF classes (**D**). Cell values give pairwise *R*^2^, and significance was estimated using 10,000 permutations followed by Benjamini–Hochberg correction within each classification. Diagonal cells denote comparisons of a class with itself. n.s., not significant; \*\*\**p_FDR_*<0.001.

